# Global transcriptome profiling uncovers footprints of root and shoot development in crop models barley and tomato

**DOI:** 10.1101/825430

**Authors:** Ali Ahmad Naz, Michael Schneider, Lucia Vedder, Bobby Mathew, Heiko Schoof, Jens Léon

## Abstract

Land plants establish their forms at two development hotspots- root and shoots apices. In this study, we dissected and compared the global transcriptome of these developmental zones in crop models barley and tomato. We employed a state of the art transcriptome analysis technique for deep profiling of expressed genes. This analysis resulted in highly reproducible quantitative expression profiles of 19,441 and 23,388 genes in barley and tomato, respectively. In barley, 16,834 genes were expressed both in root and shoot apices, whereas 1,081 genes were specific to root apex and 1,526 genes were active in shoot apex. With significant variations 20,154 genes were expressed in root and shoot apices of tomato of which 1,858 and 1,376 genes showed root and shoot specificities. Systematic analyses of these genes revealed distinct commonalties and divergence among the active genes for root and shoot development. A deeper insight in these data uncover tissue- and species specific genes, unique footprints of gene ontologies and divergence of auxins pathway genes in root and shoot apices of barley and tomato. These data provide a primary resource to understand intra- and inter-species genetic networks of root and shoot development as well as the evolution of genes in crop plants.

## INRODUCTION

Flowering plants are divided in two major classes-monocotyledons and dicotyledons. Major diversification of these plants underwent in the Early Cretaceous, between about 130 and 90 million years ago (Crane et al. 1995). In spite of a protracted evolutionary divergence most cultivated crops belong to these major categories. Among these, barley and tomato are considered genomic models for crop plants representing monocots and dicots, respectively. The major reasons behind this lie in their strict diploidy, pollination behavior, diversity and their relatedness with many economically important crop species. In addition, both crop species reveals characteristic differences in their development and growth habits especially in the root and shoot forms. Genomic dissection of this variation provides an opportunity to address key biological questions behind the evolutionary divergence of the members of monocots and dicots crop species.

Root and shoot are two main axes of plant development and growth. Roots are programmed in root apical meristem and part of elongation zone where the lateral root arise (Overvoorde et al. 2010). Shoots develop in the shoot apical meristem and its peripheral zone where leaf primordial arise successively. In the axils of these leaf primodia axillary meristems arise which give rise to axillary shoots (Naz et al. 2013). Although, root and shoot develop and grow at different locations but there exists an active communication between both organs which determines specific plant architecture. Inter-connected hormonal circuits dominated by auxin and cytokinins play a fundamental role for a coordinated development of these organs (Puig et al. 2012). There exists root-wards auxin flow from the shoot and shoot-wards flow of cytokinins from the root (Ko et al. 2014). The auxin transport from shoot to root activates strigolactones in roots which move upwards via xylem and suppresses axillary shoot branching (Puig et al. 2012). Notably, these processes are modulated by a complex but an orderly network of genes where the expressions and interplay of these genes play the pivotal role for the coordinated development and growth of plants. Notwithstanding of excellent genetic revelations and knowledge in plant development biology, it remained largely enigmatic, how and what level genotypes (having same genome) recruit different genes in establishing two contrasting growth axes? To address this, in-depth global transcriptome profiling of genes active in root and shoot and their comparison in different species was essential to gauge developmental dynamics and the evolutionary divergence of fundamental biological process in crops plants.

In the present study, we performed a global transcriptome profiling of plant development genes in root and shoot using a state of the art method-Massive Analysis of cDNA Ends (Rotter et al. 2008). This method of transcriptome analysis is known for its ultra-deep and un-biased transcriptome profiling of expressed genes from rare to the most abundant. This manuscript brought up the first report on global transcriptome profiles of genes active in root and shoots development zones. Here, we are also presenting a systematic comparison of gene transcripts based on tissue- and species specificities as well as of quantitative expression profiles of individual genes in two important crop plants.

## RESULTS

In the present study we performed in-depth transcriptome profiling of plant development gene in two crop models, barley and tomato. For this, we have dissected the root and shoot apices precisely under the microscope which represent the most relevant organs for plant development. Transcriptome analysis using MACE revealed 7.9 million reads in barley root apices of which 5.5 million reads were mapped across the barley genome. Among the mapped reads, 4.7 million reads were aligned to barley annotated genes. Relatively more transcripts (10.5 millions) were found in barley shoot of which 5.9 million reads were mapped and aligned with barley annotated genes (Figure S1A). In tomato root and shoot apices around 12.4 and 7.9 million reads were identified of which 7.1 and 5.5 million reads were mapped to annotated genes, respectively (Figure S1B). This read mapping resulted in transcriptome profiles of 17,915 genes in root apices and 18,360 genes in shoot apices of barley (Figure 1A and 1B). Comparatively more numbers of expressed genes were identified in tomato root (22,013) and shoot (21,531) apices, respectively (Figure 1C and 1D). The expressed genes revealed a very high reproducibility (correlation coefficient range from 0.97 to 1.00) among the individual biological replicates of root apices and shoot apices both in barley and tomato (Figure S2 and S3).

**Figure 1.**
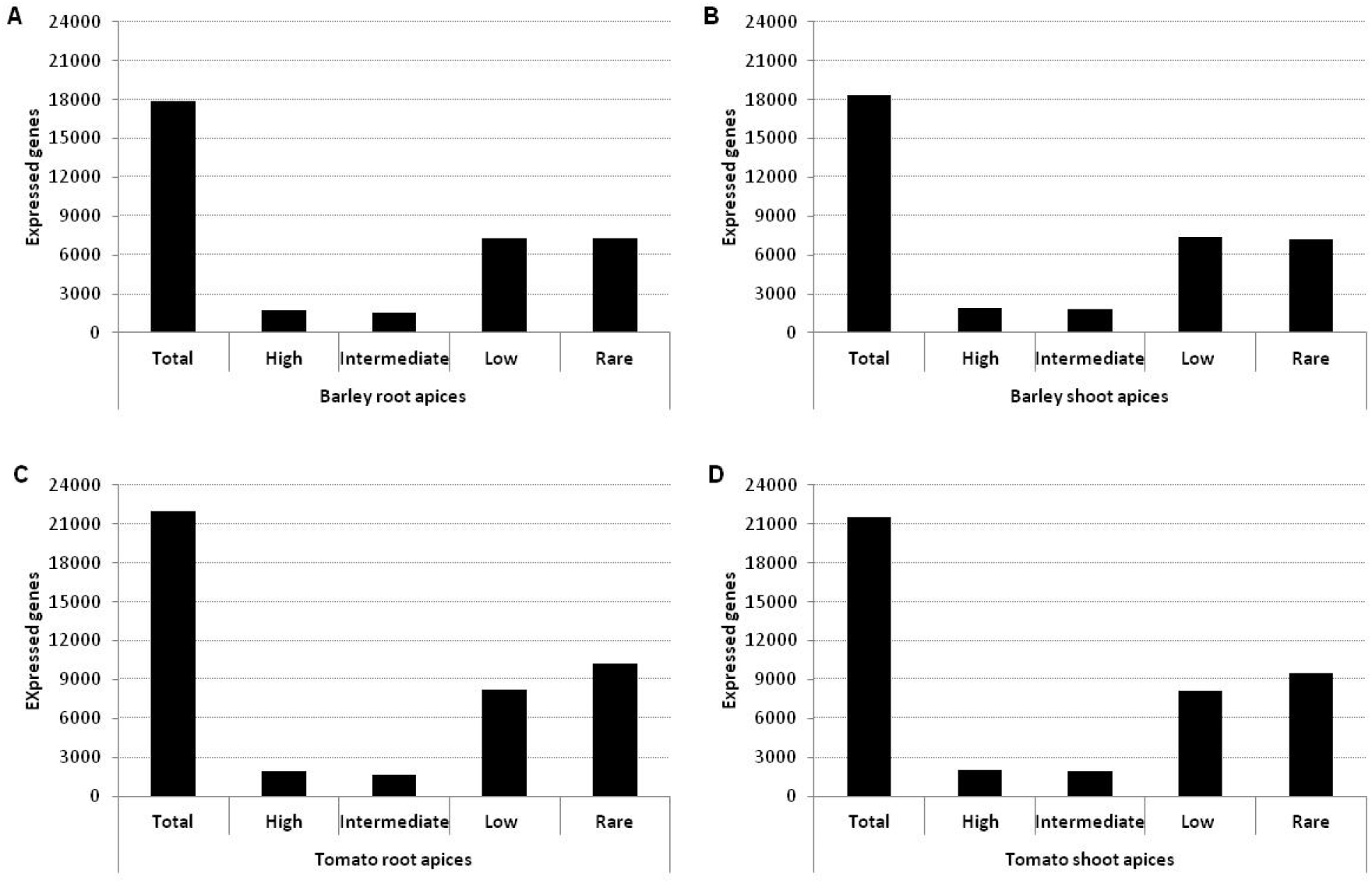
Number of expressed genes and their expression levels in barley root apices (A) barley shoot apices (B) tomato root apices (C) and tomato shoot apices (D). The levels of expression refers to rare (<5 transcripts per gene), low (>5 to <50 transcripts per gene), intermediate (>50 to <100 transcripts per gene) and abundant (>100 transcripts per gene).

To quantify the transcriptome profiles of individual genes, we categorized the expressed genes into four arbitrary classes; rare (<5 transcripts per gene), low (>5 to <50 transcripts per gene), intermediate (>50 to <100 transcripts per gene) and abundant (>100 transcripts per gene). This analysis found the substantially higher number of genes ranging in rare and low expression categories in root and shoot apices of barley and tomato.

Next we compared the expressed genes between root and shoot apices to test the commonalities and divergence of identified genes in barley and tomato. The volcano plots indicated significant set of expressed gene between root apices and shoot apices in barley and tomato (Figure S4A-B). Differential expression analysis found 1,081 and 1,526 roots and shoots apices specific genes in barley. A total of 16,834 genes were commonly expressed in root and shoot apices of barley (Figure 2A, Table S1). To link the expressed genes among different expression levels, we performed a pair-wise quantification of common genes in barley root and shoot apices (Figure 2B). Overall, common genes were identified between all categories. The highest number of common genes were identified between the category low (5,114) and rare (4,407) between the root and shoot apices in barley. Notably, there were 1,500 and 1,041 rare transcripts expressed specifically to shoot and root apices in barley. Likewise, 40 and 26 genes were specific to root and shoot apices at low level, respectively. Similar analysis in tomato identified 1,858 and 1,376 expressed genes specifically in root and shoot apices, respectively. Whereas, a major proportion of identified genes (20,154) were expressed commonly in root and shoot apices of tomato (Figure 2C, Table S2). A pair-wise comparison found the highest numbers of common genes between the root and shoot apices at the levels low and rare transcripts in tomato (Figure 2D). Similar to barley, a considerable number of rare transcripts were specific to root (1,750 genes) and shoot (1,349 genes) apices in tomato.

**Figure 2.**
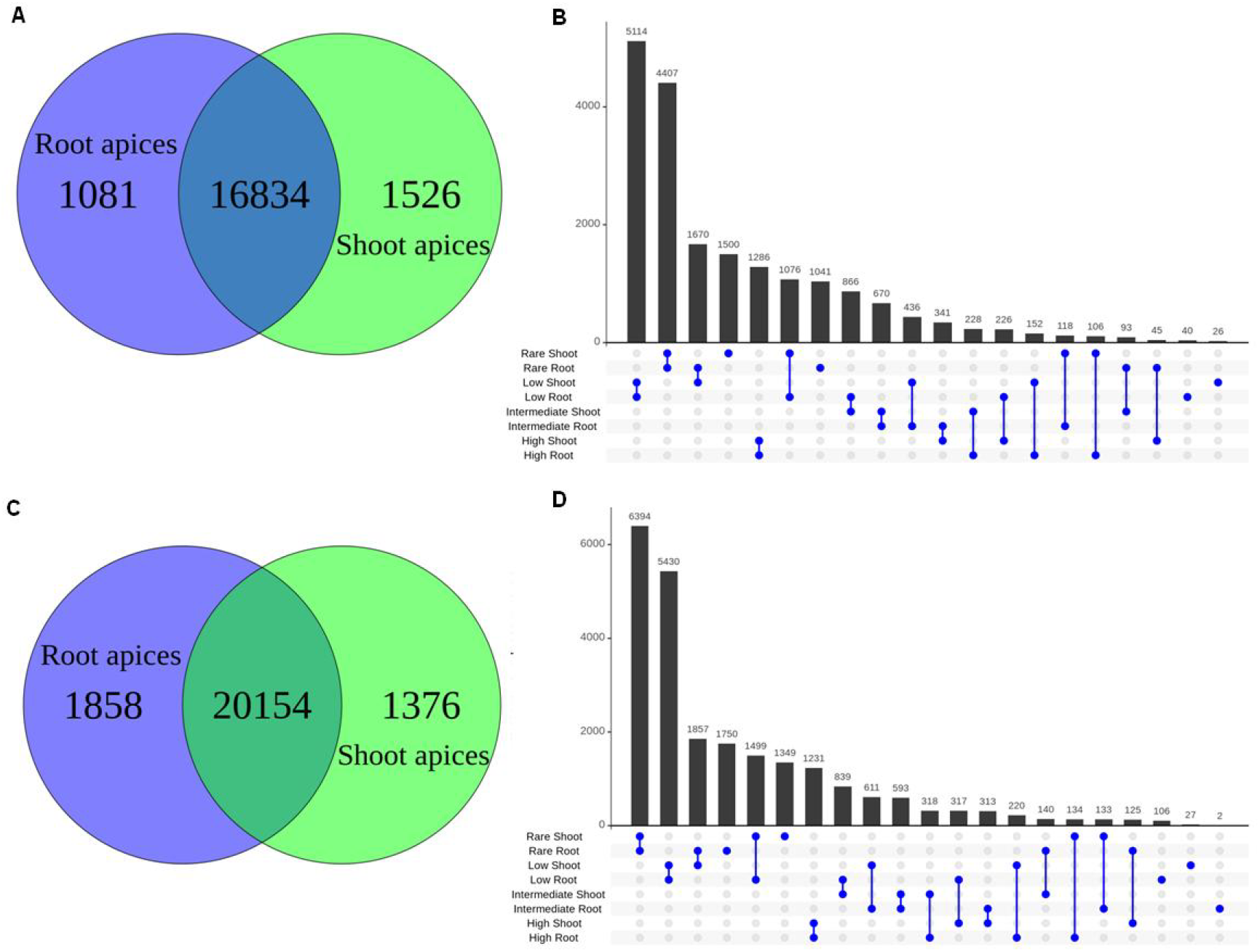
Comparison of expressed genes between barley root and shoot apices (A-B) and between tomato root and shoot apices (C-D). Different categories of expression levels refers to expression refers to rare (<5 transcripts per gene), low (>5 to <50 transcripts per gene), intermediate (>50 to <100 transcripts per gene) and abundant (>100 transcripts per gene).

In addition to the root and shoot specificities, we quantify the divergence of genes expressed commonly in root and shoot apices of barley and tomato apices. For this, first we visualized global expression profiles of individual transcripts in two separate circos plots across the barley genome revealing normalized expression for three biological replicates of root and shoot apices (Figure S5 and S6). Later, we log_10_ transformed the transcript data in counts per millions (CPM+1) scale where expression ranges from 0 (not expressed) up to 4 (highly expressed), and plotted heat maps between the root and shoot apices of barley and tomato. In barley, among 16,834 genes which were expressed both in root and shoot, 1,491 (8.8%) revealed significant variation in root and shoot apices (Figure 3A). Remarkable expression patterns were clustered in four main groups. Similarly, we compared the 20,154 genes expressed in root and shoot apices of tomato. This analysis found 1,990 (9,9%) genes that revealed significant variation between the root and shoot apices of tomato (Figure 3B). Overall, both species showed a comparable gene clustering based on the differential expression variation. Altogether, we found around 4,098 genes which are likely to be differential expressed in root and shoot apices of barley and 5,224 potentially differential expressed genes in root and shoot apices of tomato.

**Figure 3.**
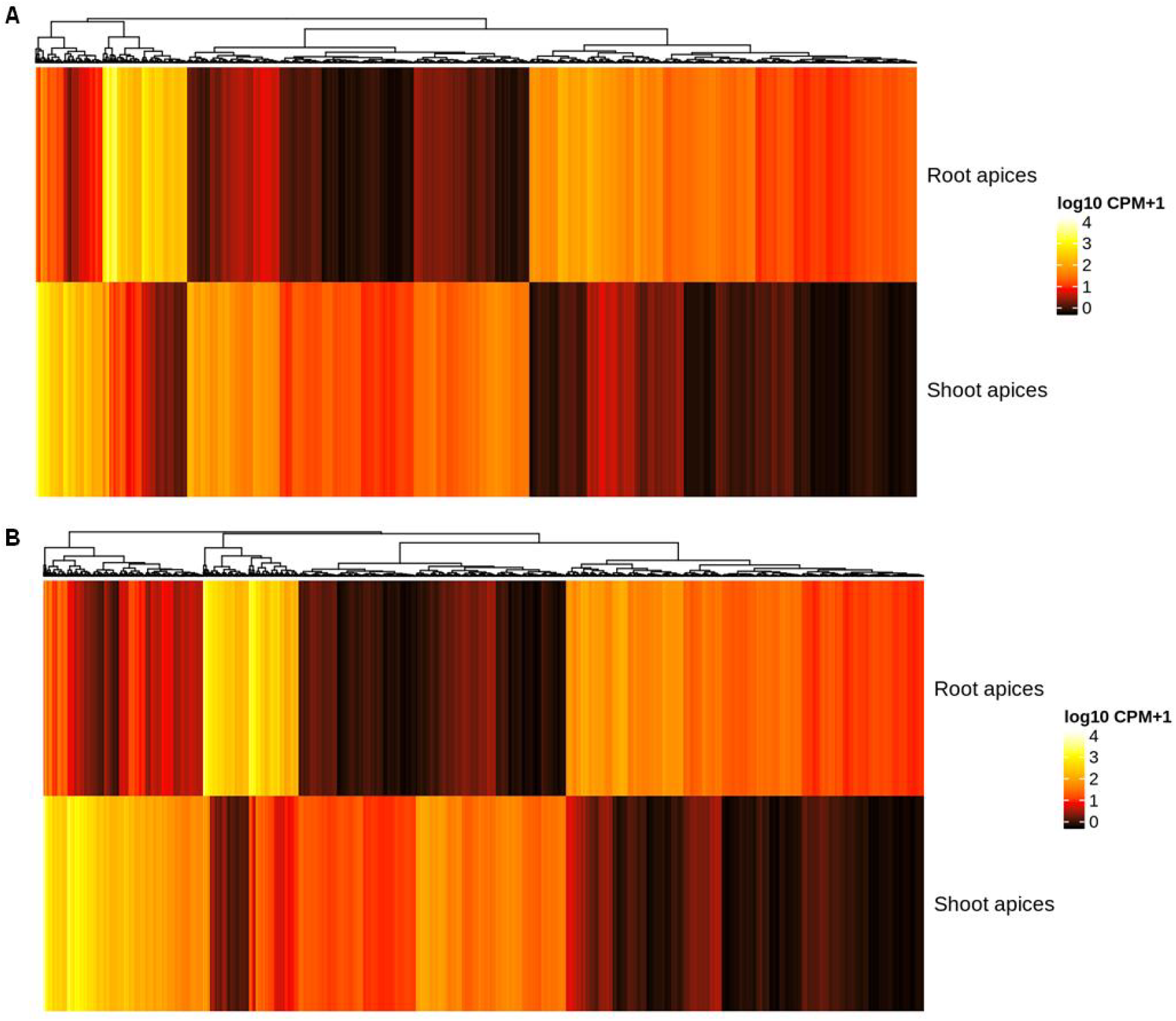
Differential expression of genes between barley root apices and shoot apices (A) and between tomato root apices and shoot apices (B). The scale (0 to 4) represents log10 counts per millions (CPM+1) of gene transcripts. Scale: 0 (not expressed) up to 4 (highly expressed).

Next we classified the differentially expressed gene in root and shoot apices based on the gene ontology (GO) terms. Among the differentially expressed genes, the highest number of genes expressed in barley root belongs to GO:0055114 (oxidation-reduction process) and GO:0020037 (heme-binding process). In barley shoot apices, a predominant number of expressed gene were the DNA binding transcription factor proteins (GO:0003677) (Figure 4A, Table S3). Similarly, the highest numbers of gene revealing differential expression were found on tomato root apices which belong to perioxidase activity (GO:0004601). In tomato shoot apices, the highest number of differential expressed were related to photosynthesis process (GO:0015979) (Figure 4B, Table S4). As phytohormone auxin play a primary role in the development and growth, we visualize the expression divergence two related gene families, Auxin-responsive proteins (Indole-3-Acetic Acid, IAA) and auxin response SAUR-proteins (Small Auxin Up RNA) involved in the auxins pathway. The member of these gene families were more (in number) and highly expressed in root apices as compared to shoot apices both in barley and tomato (Figure S7 and S8).

**Figure 4.**
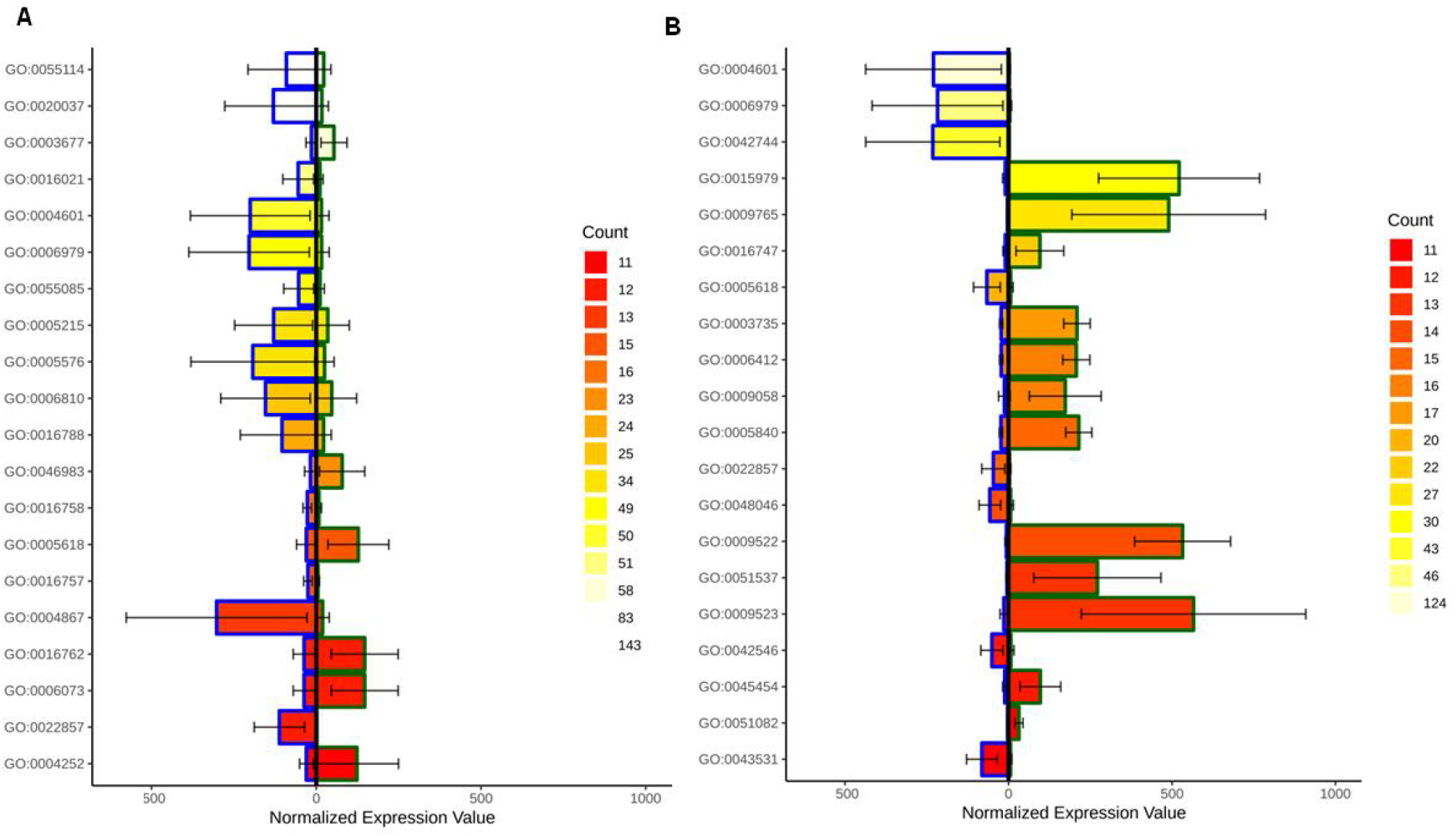
Gene ontology enrichment analysis of differentially expressed genes between barley root apices (left) and shoot apices (right) (A) and between tomato root apices (left) and shoot apices (right) (B). Scale represents gene counts in numbers and error bars showed standard deviation of normalized gene expression in each category.

Finally, gene expression analyses were performed between tomato and barley root and shoot apices. For this, initially true orthologous genes of barley and tomato were identified using OrthoMCL gene clustering. This analysis found a set of 519 genes, showing a 1-to-1 relationship between barley and tomato which were depicted across the barley and tomato genomes (Figure 5, Table S5). This 1-to-1 relationship means that the respective barley and tomato gene are more closely related one to each other than to every other gene.

**Figure 5.**
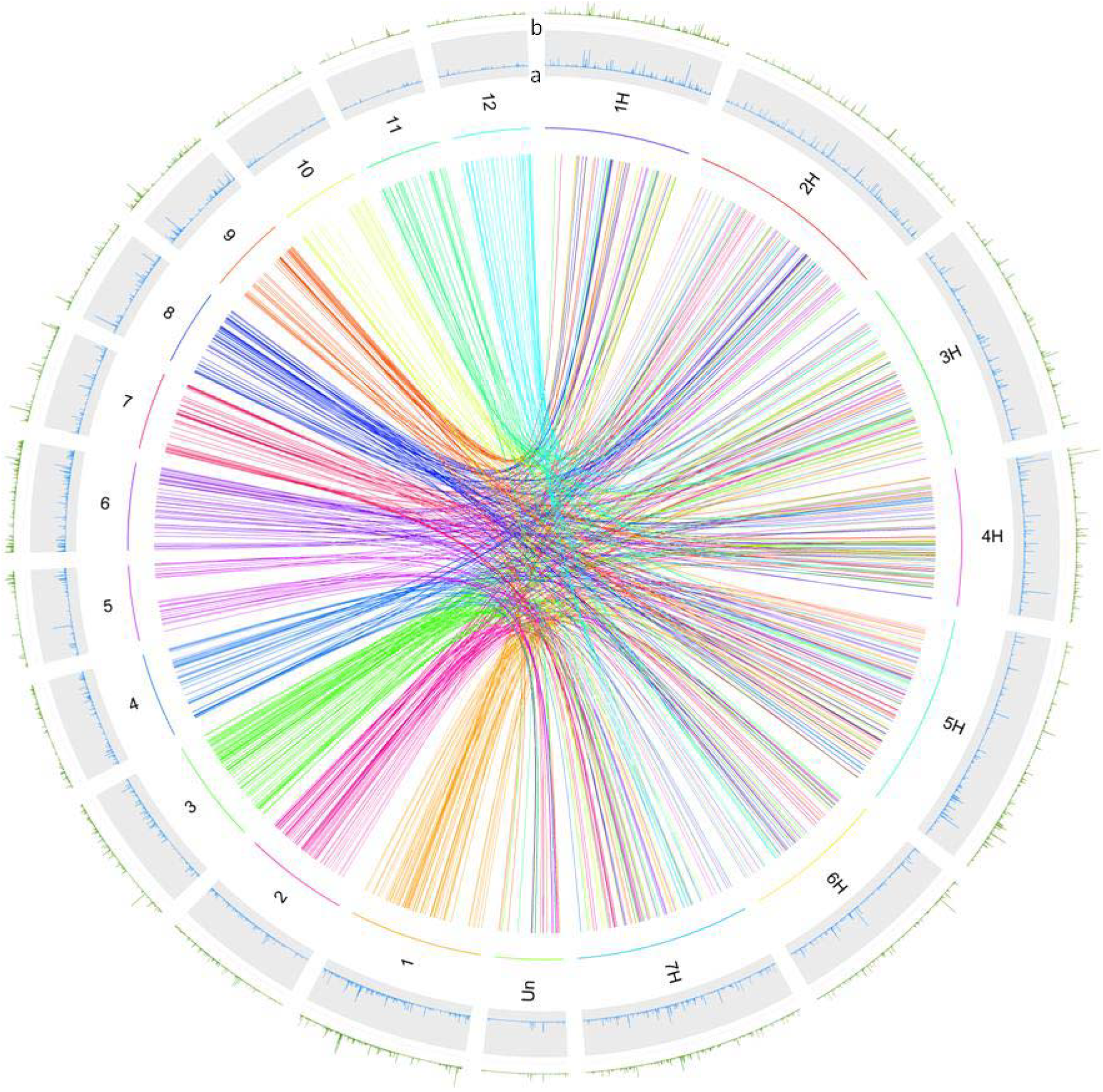
Circos-plot of barley and tomato genomes showing the position of orthologous genes having a 1-to-1 relationship. The chromosomes 1H to 7H and 1 to 12 represent the chromosomes of the barley and tomato genome, respectively. The colors of the links are related to the twelve tomato chromosomes for better visualization. The position of othologous genes between barley and tomato can be read following the link bands in the center. The outer two circles show the mean expression for root apices (blue, a) and shoot apices (green, b).

Differential expression of the selected 519 orthologous genes of barley and tomato between root and shoot apices is presented in a multi comparison heatmap (Figure 6A). This heatmap is created using R tools Complex Heatmap that showed log_10_ CPM+1 scale where expression ranges from 0 (not expressed) up to 4 (highly expressed). For this, the libraries where tested on a global scale for the identified 519 orthologous genes against each other using a generalized linear model based on a binomial distribution. These values indicate a gradual difference of the barley root apices library, following the arithmetic same species < same tissue < different tissue. The overlap of barley apices is higher to the tomato root apices compared to the tomato shoot apices. For comparison, both sets of barley and tomato for root and shoot apices were merged into one data source, because no gene size error has been introduced using a CPM normalization. On a given threshold (p-value 0.05), 138 differential expressed genes were identified between the tissues, whereas 209 genes revealed significant variation between species. Differential expression analysis between tomato and barley on single gene level was performed using all replicates of root apices and shoot apices. Thirteen tomato genes show up regulation compared to their barley relatives, while eight of these do not show any expression in barley at all. Contrarily, nine genes revealed up regulation in barley apices compared to tomato apices. In this group, one single gene was missing in the tomato apices, while the remaining 8 were lowly expressed. The gene groups that are tissue and not species specific were apparently smaller. Two genes related to shoot and three genes related to root were identified at p-value 0.05. For instance, the BAG chaperone regulator 7 was up regulated in tomato apices while down regulated in barley apices, whereas the BAG chaperone regulator 4 showed an opposite pattern. A similar pattern was found for glutathione s-transferase genes. An aquaporin like protein as root specific over the species levels as well as a pollen allergen as shoot specific were differentiated from the other two groups (Figure S9).

**Figure 6.**
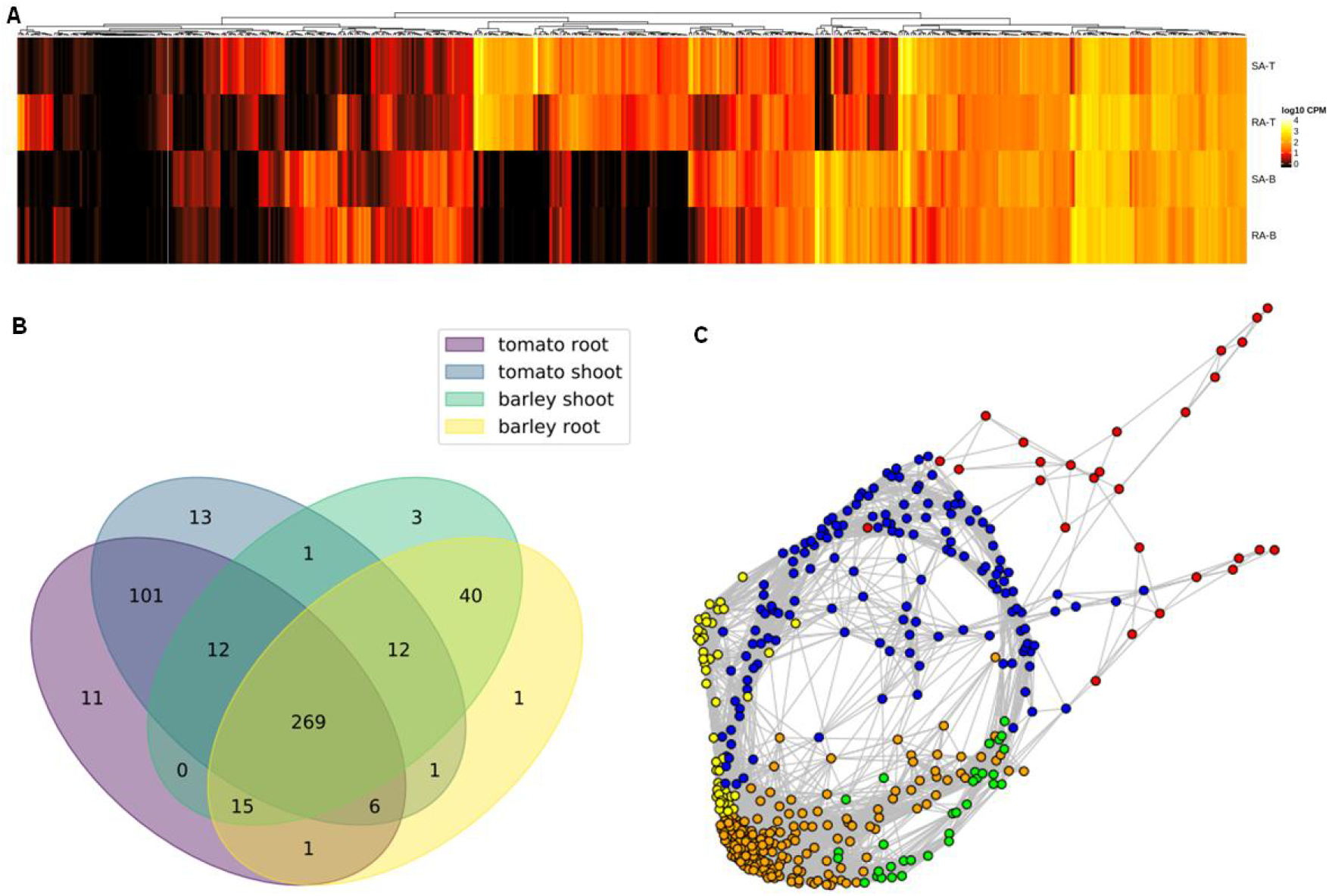
(A) Differential expression analysis of 519 orthologous genes of barley and tomato in root and shoot apices. Scale: log_10_ CPM+1 where expression ranges from 0 (not expressed) up to 4 (highly expressed). SA-T = shoot apices tomato, RA-T = root apices tomato, SA-B = shoot apices barley, RA-B = root apices barley. (B) Venn diagram showing the number of expressed genes in barley and tomato root and shoot apices. (C) Co-expression network analysis of the selected orthologous genes (cluster 1=red, cluster 2=green, cluster 3= blue, cluster 4=yellow, cluster 5=orange).

The count of differentially expressed genes species by species exceeds the count of differentially expressed genes related to a root or shoot apices regardless of the border of species. Of those 519 genes, 486 ones are expressed in at least one species and tissue (Figure 6B). Genes are marked as expressed if they have an expression value higher than 0. For each species and tissue the union of expressed genes in each replicate is used for the comparison in between species and tissues. In total, there are 415 genes expressed in tomato root apices, 415 genes expressed in tomato shoot apices, 345 genes expressed in barley root apices, and 352 genes expressed in barley shoot apices (Figure S10). A major proportion of these genes (269) were expressed in all four subsets whereas 125 and 44 genes were only expressed in tomato and barley respectively. Among these, 207 genes showed differential expression in root apices and 217 in shoot apices of barley and tomato. Testing for significance using FDR (<1%) leads to 69 genes being highly differentially expressed in root apices and 77 genes being highly differentially expressed in shoot apices, respectively (Table S6 and S7). Next, we performed a co-expression network analysis in order to investigate concerted action of orthologous genes in tomato and barley. We used graphical Lasso (Friedman et al. 2008) to construct the network and the network was subdivided using a fast-greedy community algorithm (Clauset et al. 2004). Our co-expression analysis resulted in total five clusters indicated in different colors (Figure 6C). Among these, the node with the highest numbers of co-expression links belongs to a Zinc finger CCCH domain-containing protein 2 (HORVU3Hr1G019510, Solyc12g008660). The list of the genes belonging to each cluster is provided as the supplementary material (Table S8 to S12).

## DISCUSSION

In the present study, we aimed in-depths global expression profiling and comparison of developmentally active genes in these organs in two major crop species. For this, root apices (7 mm) comprising of root meristem and root elongation zone of barley and tomato were precisely harvested under the dissecting microscope. Likewise, shoot meristems comprising of two emerging leaf primordia were harvested in both species. To homogenize the sampling process, we cut exactly 50 root and shoot apices (as technical replicates) and pooled them in each biological replicate. The primary reason behind this sampling strategy was to target development related genes and to ensure reproducibility of transcript data. Our data showed very high reproducibility among the individual biological replicates in each tissue in both species suggesting that the adopted sampling strategy was appropriate. The present transcriptome profiling was made using a state of the art method-Massive Analysis of cDNA Ends (MACE). This method is based on the TrueQuant approach that recognizes PCR-bias and eliminates it during the transcriptome sequencing. In TrueQuant, PCR-template molecules are labeled with the unique TrueQuant Adapter or “UMI” prior to their amplification, such that each molecule consists of a unique sequence (GeneXPro, Frankfurt, Germany). By this, ideally each template molecule can then be identified by its unique TrueQuant adapter sequence. Based on this, PCR copies can be identified and eliminated from the dataset and hence uneven amplification and artifacts generated during the PCR amplification can be eliminated (Rotter et al. 2008). In addition, MACE target only one single read per transcript molecule is sequenced, short and rare transcripts are identified already at 10-20 times lower sequencing depth, when compared to full length RNA-Seq and no length-based normalization is required (Zawada et al. 2014, (Soneson and Delorenzi, 2013). Therefore, the employed method in this study was suitable for ultra-deep and un-bias transcriptome profiling and seems advantageous to standard whole genome RNA sequencing techniques, which rely on random hexamer primers for amplification (Hansen, Brenner and Dudoit, 2010).

Our analyses uncover a total of 17,915 and 18,360 genes in barley root and shoot apices, respectively. Higher numbers of genes were expressed in tomato root (22,013) and shoot (21,531) apices. A primary reason behind this may lie in comparatively improved genome assembly and gene annotation in tomato as compared to barley. Evolutionary, a putative reason may lie in extreme genome divergence between both species. Barley carried extremely longer 5.3 Gb of genome sequence as compared to 828 Mb in tomato (Mascher *et al.*, 2017), The Tomato Genome Consortium, 2012). The whole genome sequencing in barley and related species like wheat has found the presence of high proportion of repetitive elements and their higher mutation rates (International Wheat Genome Sequencing Consortium (IWGSC), 2018, Luo et al. 2017). Such characteristics of genome evolution may have a role in pseudogenisation resulting in a slightly lower number of expressed genes in barley as compared to tomato. A comparable transcriptome study in barley found a total of 25,152 transcripts expressed in leaf and main shoot meristem tissues of which 4,044 and 2,340 were exclusively expressed in leaf and shoot apices, respectively (Digel et al. 2015).

Next we categorized gene expression of root and shoot apices in four expression levels; rare, low, intermediate and high. Although this classification was arbitrary, it greatly helped to quantify the gene expression divergence or commonality between different tissues in each species. We found that the numbers of rare and low levels transcripts were predominantly high in root and shoot apices in barley and tomato. Also, the highest numbers of gene expressed commonly both in root and shoot apices of barley and tomato were fallen in rare and low levels transcripts categories indicating the significance of these genes in plant development. (Feng et al. 2017) performed a similar expression quantification of the expressed genes in wheat inflorescence. They have found the prevalent number of lowly expressed genes in wheat reproductive meristems. The comparison of expression divergence between tissues showed significant variation in root and shoot specificities, these variation was comparable in barley and tomato suggesting an overall gene conservation pattern in barley and tomato. These results are in line with (Calixto et al. 2015) in which they study evolutionary relationship of Arabidopsis core circadian clock and clock-associated genes with several member of monocot and dicot species. They found high gene conservation of orthologous and paralogous genes in different species and variation in gene copy numbers like gene duplications among the various species. To see this pattern on individual gene levels, we performed OrthoMCL gene clustering to test 1-to-1 relationship of expressed genes in barley and tomato. Here, we found that a major portion of 159 genes were expressed in all four subsets, 74 genes expressed only in tomato and 59 genes that were expressed in barley which can be promising candidate for evolutionary studies on barley and tomato. More number of root and shoot specific genes were found in tomato than barley which correspond with the total genes identified in both species. Overall, these data again suggest higher gene conservation between the species than the selected plant development hotspots-root and shoot apices. However, cumulative gene analysis using gene ontology identified unique pattern and prevalent gene numbers involved in oxidation-reduction and heme-binding processes in barley root whereas perioxidase activity genes in tomato root apices. Likewise, remarkable differences were observed among the auxin-responsive proteins and auxin response SAUR-proteins involved in the auxins pathway. More number and higher expression of the member of these gene families were found in root apices as compared to shoot both in barley and tomato suggesting more auxin activity in root as compared to shoot. This notion support the shoot to root auxins transport that seems critical for a balanced and coordinated development of roots and shoots (Ko et al. 2014, McDavid et al. 1972).

Take together, the present data provide a first comprehensive transcriptome profiling of root and shoot development genes (from rare to abundant) and their systematic comparison in two crop model species. Here, we presented quantitative expression map of 17,915 (in root) and 18,360 (in shoot) of active gene in barley. Similarly, expressional divergence of 22,013 and 21,531 has shown in tomato root and shoot apices, respectively. Comparative analyses uncover unique conservation as well as divergence of gene expression between the developmental hotspots of barley and tomato. These data have also shown its utility to study individual gene as well as cumulative analysis of genes related to a given pathway or biological process at tissue and specie specificities levels. We believe, these data will act as new resources which greatly complement the genomic knowledge of plant development in crops. Further, it will help to extending future research on plant developmental and evolutionary biology of crop plants.

## Materials and Methods

### Plant material and experimental design for transcriptome analysis

In the present study, we utilized spring barley cultivar Scarlett and tomato cultivar Moneymaker. The seeds of respective genotypes were pre-germinated and sown in soil in 96-cell plant growing trays on consecutive dates. The plant was grown inside the growth chamber adjusted at 22°C for 8 hours (in the day) and 18°C for 16 hours (at night). The plants of each 96-cell tray were grown under these conditions for 10 days. After this, root and shoot apices were harvested from one 96-cell trays as one biological replicate. For root apices, 7 mm (containing apical meristem and elongation zone) of the primary roots of barley and tomato were harvested independently and a total of 50 root apices were pooled in each biological replicate. Likewise, three biological replicates were harvested independently in each species. Barley vegetative shoot apex comprising of apical meristem and emerging leaf primordia. Here again, we harvested 50 shoot apices and pooled them in each biological replicate. Similarly, 50 tomato vegetative shoot apices comprising shoot apical meristem along with two emerging leaf primordial were collected. The harvested samples were sent to GeneXpro GmbH (Frankfurt, Germany) for transcriptome analysis. Transcritome analysis was made in three biological replicates of root (having 50 technical replicates) and shoot (having 50 technical replicates) apices in both species using Massive Analysis of cDNA Ends (MACE) by TrueQuant method according (Rotter et al. 2008).

### Gene Expression Analysis

#### Upstream analysis profile

Sequencing MACE data derived from GeneXPro GmbH was quality trimmed using Trimmomatic SE (version 0.36) using a minimum length setting of 40 bases and moderate quality parameters 28 for the leading and 17 for the trailing bases linked with a headcrop of 10 bases (Bolger et al. 2014). Fragments were aligned with the barley (IBSC_V2) and tomato (SL2.50) reference genome using bwa (version 0.7.17) applying standard settings. Read filtering was performed strictly, applying a quality filter of >60 using samtools 1.8 view option (Li, 2011). Duplicates as residuals from the PCR step in sequencing where not regarded due to the low impact of these in expression analysis reported by Parekh et al. 2016 (Parekh et al. 2016) and the applied TrueQuant technique patented by GenXPro to remove most of the PCR duplicates from sequencing. The fragments were matched to the genes by the tool feature counts of the subread software package (version 1.6.2) for tomato and barley separately, using the corresponding annotation files for the used reference (Liao et al. 2013).

#### Downstream analysis profile

Further analysis was performed in the R (3.4.4) and Julia (1.0.3) environment. Expression comparison and plotting was based on counts per million (CPM) normalized values.

Replicate testing was done applying a generalized linear mixed model based on a negative binomial distribution and correlation analysis. Later on, genes are clustered in four groups of expression levels – rarely expressed genes showing expression with less than 5 CPM fragments, low expressed genes with >5 to <50 CPM fragments, intermediate expressed genes with >50 to >100 CPM fragments and highly expressed genes having a higher expression than 100 CPM fragments.

The intra-specific differential expression analysis was performed based on adjusted parametric test on all replicates of root against shoot apices and the log fold-change.

To go further on and be able to compare barley to tomato expression on genome level, potential orthologous genes with a family size of one are searched based on a clear 1-to-1 relationship between the genes of barley and tomato. This task was performed by using the OrthoMCL tool (version 2.0.9) to cluster the reference peptide sequences of tomato (SL2.50) and barley (IBSC_v2).

The information obtained from the gene orthologous analysis was linked to the expression data and gene function and GO term information. Functional annotation was taken from the IPK barley IBSC project. Mean expression values over the replicates where clustered and plotted.

The libraries of barley and tomato root and shoot apices were tested against each other as whole groups, consisting of the mean expression of selected 519 orthologous genes. Potential variations on the library level were identified using a generalized linear model based on a negative binomial distribution.

Potential differential expressed genes in barley to tomato tissue for the orthologous genes were further analyzed on single gene level to reveal the impact of single genes in this subgroup. Therefore, all six replicates of root and shoot apices were merged and tested in a generalized linear model based on a negative binomial distribution against the other group for significant differences. The same procedure was followed for species expression variation of barley and tomato.

Venn diagrams were prepared using R package VennDiagram 1.6.20 (Hanbo, 2018) and UpSetR 1.3.3 (Nils, 2017), circos plots by the R package OmicCircos 1.16.0 (Ying, 2015). Bioconductor package ComplexHeatmap 1.17.1 was used to create the heatmaps, Correlations and Go term plots of the replicates were printed using Julia package Gadfly v1.0.1 (Gu et al. 2016).

Gene co-expression were made using graphical Lasso (Friedman et al. 2008) to construct the network and the network was subdivided using a fast-greedy community algorithm (Clauset et al. 2004).

## Author’s Contribution

AAN and JL conceptualize the research. MS, AAN, BM, LV, HS analyze the data. AAN, MS and LV have written the manuscript.

## Acknowledgements

We are grateful to Mrs. A. Bungartz for her help in sample preparation. Special thanks to Mrs. Anna Vlasova for her valuable suggestion in data analysis and to Mr. Md. Kamruzamman for reading the manuscript.

